# Adaptive multipole models of OPM data

**DOI:** 10.1101/2023.09.11.557150

**Authors:** Tim M Tierney, Zelekha Seedat, Kelly St. Pier, Stephanie Mellor, Gareth R Barnes

## Abstract

Multipole expansions have been used extensively in the Magnetoencephalography (MEG) literature for mitigating environmental interference and modelling brain signal. However, their application to Optically Pumped Magnetometer (OPM) data is challenging due to the wide variety of existing OPM sensor and array designs. We therefore explore how such multipole models can be adapted to provide stable models of brain signal and interference across OPM systems. Firstly, we demonstrate how prolate spheroidal (rather than spherical) harmonics can provide a compact representation of brain signal when sampling on the scalp surface with as few as 100 channels. We then introduce a type of orthogonal projection incorporating this basis set, the Adaptive Multipole Models (AMM), which provides robust interference rejection across systems, even in the presence of spatially structured nonlinearity errors. This projection is always stable, as it is an orthogonal projection, and will only ever decrease the white noise in the data. However, for array designs that are suboptimal for spatially separating brain signal and interference, this method can remove brain signal components. We contrast these properties with the more typically used multipole expansion, Signal Space Separation (SSS), which never reduces brain signal amplitude but is less robust to the effect of sensor nonlinearity errors on interference rejection and can increase noise in the data if the system is sub-optimally designed (as it is an oblique projection). We conclude with an empirical example utilizing AMM to maximize signal to noise ratio (SNR) for the stimulus locked neuronal response to a flickering visual checkerboard in a 128-channel OPM system and demonstrate up to 40dB software shielding in real data.

## 1. Introduction

Multipole expansions provide a model-based approach to analysing data acquired from MEG systems. These expansions most often utilise spherical harmonics and decompose the data into spatial patterns arising from the brain (internal harmonics) and from sources of magnetic interference (external harmonics). This approach, which is termed Signal Space Separation or SSS (Taulu et al., 2005; Taulu & Kajola, 2005), makes minimal assumptions about the anatomy of the participant and only requires that the sensor array geometry and calibration is known (Nurminen et al., 2008). SSS can also be expanded to remove temporally correlated sources of interference (termed TSSS, (Taulu & Simola, 2006)), which typically occur when the source of magnetic interference is near the sensor array, or calibration errors are present. In addition, these methods can be used with continuous head position monitoring to correct for motion artefacts in cryogenic MEG systems (Wehner et al., 2008).

Software based correction of motion artefacts and environmental interference is a key (and currently unresolved) issue in the development of wearable OPM systems (Brookes et al., 2021; Ding et al., 2023; Hill et al., 2022; Holmes et al., 2023; Mellor et al., 2021; Nardelli et al., 2020; Rea et al., 2021; Seymour et al., 2021, 2022; Tierney et al., 2021; Wang et al., 2023). Thankfully, multipole models of magnetic field are uniquely suited for suppressing the interference encountered in this scenario (Tierney et al., 2022). This is because, when working with on-scalp OPM sensors, the model of brain signal and interference span the same space regardless of head position or orientation (when the head does not move with respect to the sensors). This simple property of on-scalp MEG (regardless of sensor type) enables the use of these models during participant movement without any head tracking. This is qualitatively distinct from Superconducting Quantum Interference Device (SQUID) based MEG where head-tracking is required for motion correction (Stolk et al., 2013; Wehner et al., 2008). This situation also contrasts with methods based on reference arrays or decompositions of empty room noise recordings which assume a fixed (or known) spatial relationship between reference array and sensor array (Fife et al., 1999; Robinson et al., 2022; Uusitalo & Ilmoniemi, 1997; Vrba & Robinson, 2002).

To date, the use of multipole models with OPM data have largely been limited to modelling the external interference space and neglecting the models of the neuronal space (Mellor et al., 2021; Rea et al., 2021; Tierney, et al., 2021). There are a number of reasons that have, thus far, limited the use of these multipole models of neuronal signals in OPM data. Most notably, the number of channels required needs to be greater than the number of harmonics modelled. Our recent work suggested that a minimum of 150 channels would be required (Tierney et al., 2022). Over time larger OPM arrays have been created and this issue has become less of a concern (Alem et al., 2023; Boto et al., 2022; Pratt et al., 2021). However, a more fundamental issue is one of convergence. The application of these harmonic models only guarantees convergence when a single sphere that contains the whole brain can be placed fully inside the sensor array, with no sensors outside of the sphere. For MEG data sampled at the scalp surface, there may not be an origin that can be selected to meet this criterion (Zhdanov et al., 2023). As a result, the convergence of a spherical harmonic model is suboptimal, most notably affecting areas at the front and back of the head (Tierney et al., 2022).

Another issue is one of stability. The sensitivity of multipole modelling to noise (small perturbations in the signal) is determined by the condition number of the multipole model (Nurminen et al., 2010; Taulu & Kajola, 2005) and the condition number is highly dependent on array (and sensor) design (Nurminen et al., 2013). For instance, purely radial or purely tangential sensor arrays will not result in a stable multipole model (Nurminen et al., 2010; Tierney et al., 2022). As such, careful optimisation of the model’s harmonic order or substantial numerical regularisation may be required (Wang et al., 2023).

Here we propose adaptations to our multipole model to address these issues of convergence and stability. In the literature on gravitational modelling, it is common to overcome issues of non-convergence/divergence with the use of ellipsoidal or spheroidal harmonics as opposed to spherical harmonics (Fukushima, 2014; Hirt & Kuhn, 2017; Hu, 2012; Levie, 1971; Lowes & Winch, 2012; Maus, 2010; Reimond & Baur, 2016; Šprlák & Han, 2021; Winch, 1967). We propose a similar solution for on-scalp MEG data. As the geometry of the brain is well described by a prolate spheroid (the anterior-posterior axis longer than the other two), we will model signals from OPM recordings using a solution to Laplace’s equation in prolate spheroidal coordinates.

In terms of stability, we have previously taken the approach of orthogonalizing both our lead fields (brain signal) and data with respect to our model of interference (Tierney et al., 2021). This approach is inherently stable, regardless of array design, because the projection is an orthogonal projection. Orthogonal projections guarantee that the variance of the projected data will be less than the variance of the original data (see Supplementary text S1 for proof). In contrast, SSS is an oblique projection (which does not have this property) and necessitates the MEG system is designed suitably so as to maximise stability.

In principle, these adaptations should make our multipole models more stable and geometrically adaptive across different array designs. We will therefore examine the convergence properties, interference rejection abilities, susceptibility to nonlinearity errors and stability of such a multipole model. We will then conclude with an empirical example of these AMMs being used to analyse real OPM data.

## 2. Theory

### 2.1 Spherical and Spheroidal harmonics

In a space with no free current, the magnetic field (***B***) can be represented as the gradient of the magnetic scalar potential (*V*)

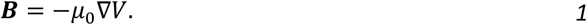

This potential can be expanded in terms of spherical harmonics (*Y*_*lm*_), using spherical coordinates (*r, θ, φ*) as follows,

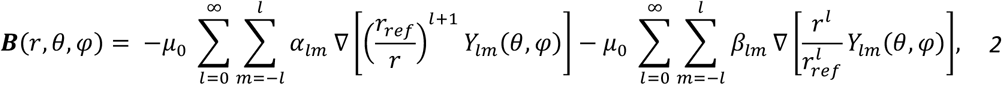

Where

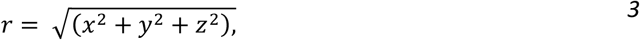

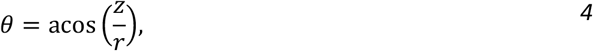

and

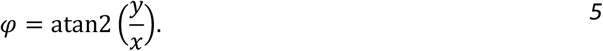

The order variable (*l*) is typically used to truncate the series at the point where it is considered to have converged (e.g. 8 for the first term in the sum and 3 for the second term). The variables *x, y* and *z* are the standard Cartesian coordinates. The spherical coordinates *r, θ, φ* are the radius, co-latitude and longitude respectively. The constant *r*_*ref*_ is the radius of a minimum volume reference sphere encasing the brain. As this term is a constant, it is rarely included in the MEG literature but is common in the gravitational literature (Reimond & Baur, 2016). It is also useful to help one gain an intuitive understanding of the series convergence. As the harmonic order (*l*) tends to infinity, the first term tends to zero for coordinates outside the reference sphere radius (*r* > *r*_*ref*_), meaning that this term models signals originating from within the reference sphere (i.e. brain signal). In contrast, as the harmonic order (*l*) tends to infinity, the second term tends to zero for coordinates inside the reference sphere radius (*r* < *r*_*ref*_), meaning that this term models signal originating from outside the reference sphere (interference).

We now examine the solution to Laplace’s equation in prolate spheroidal coordinates. This solution differs to the spherical case in that the radial dependence is replaced with ratios of the associated Legendre functions of the first 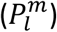 and second kind 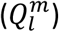.

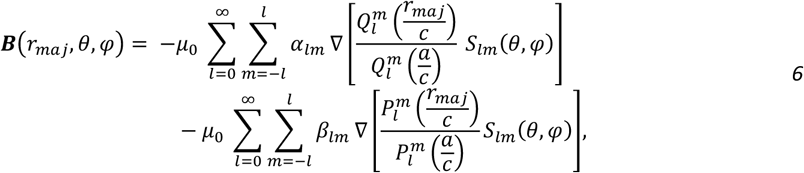

The ratios of these associated Legendre functions are provided below (Abramowitz & Stegun, 1965; Winch, 1967)

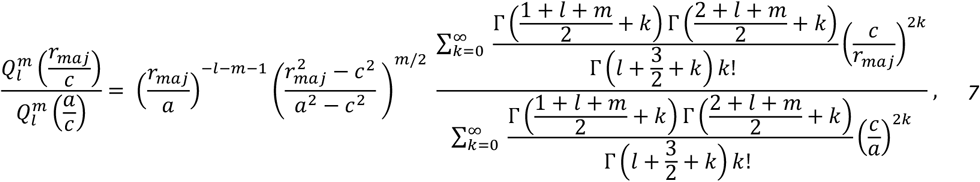

and

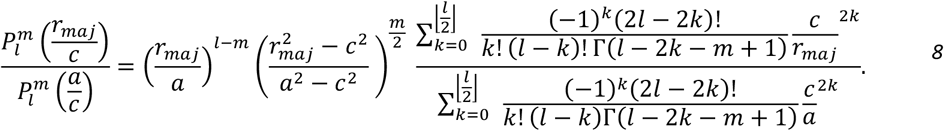

The above functions introduce the major axis coordinate (*r*_*maj*_), as well as constants (*a, b* and *c*) which define the reference spheroid enclosing the brain (See Figure 1A). These constants refer to the major axis, minor axis and focus (*a, b, c*, respectively) of the reference spheroid. The gamma symbol (Γ) refers to the gamma function. Note the longitude and latitude prolate spheroidal coordinates (*θ, φ*) have a similar interpretation to those used in the spherical case. We also change from using the complex harmonics (*Y*_*lm*_) to the real harmonics 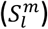 which are more interpretable when modelling real magnetic fields

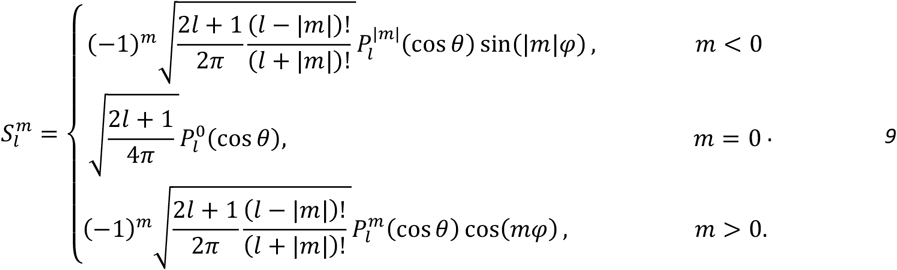

**Figure 1.**
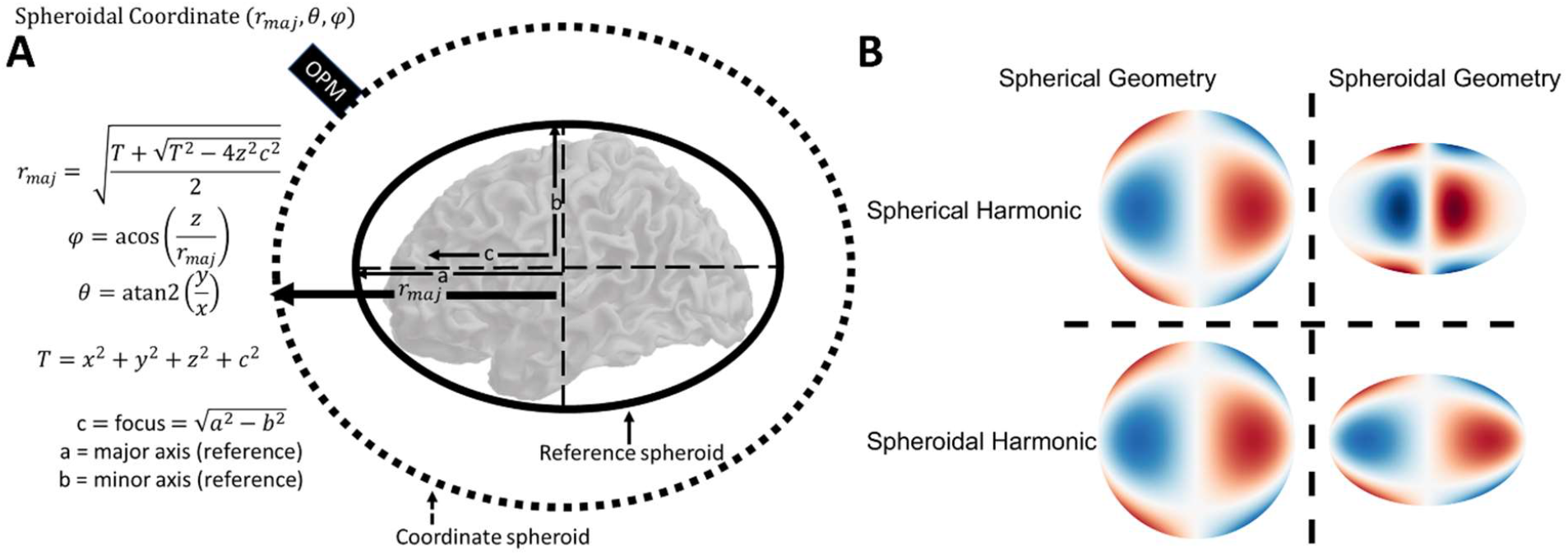
Spheroidal harmonic coordinate system and example harmonics. In A, the coordinate system of the prolate spheroidal harmonics is described. Initially, a reference spheroid is fit to the brain (or sensor array) parameterised by a reference major axis (a) and a reference minor axis (b). Once a reference spheroid is defined the coordinate of the sensor can be defined in prolate spheroidal coordinates (r_maj_, θ, φ). In B we provide an illustrative example of both spherical and spheroidal harmonics (order: l=3, m=2) on a sphere and spheroid. Importantly, when spherical or spheroidal harmonics are evaluated on a sphere they produce identical results (as the sphere is a spheroid with axes all equal in length). However, when the geometry is spheroidal (right column) we notice that the spherical harmonic model incorrectly predicts most signal to be close to the coordinate system origin and very little along the major axis (left-right direction in this figure).

It is important to note that while the spheroidal harmonics are more mathematically complicated than the spherical harmonics, they are proportional to the spherical harmonics when either the focus is zero (*c* = 0) and as such the geometry of the reference object is modelled by a sphere, or the major axis coordinate is much larger than the focal distance (*r*_*maj*_ ≫ *c*). The second scenario occurs when the sensors are far from the brain/scalp (cryogenic MEG systems). When the geometry of the object becomes more spheroidal, the spherical model predicts less signal along the major axis and most signal is predicted close to the coordinate system origin. In contrast, the spheroidal model predicts a more even signal distribution across the major axis dimension. As such, the spheroidal harmonics can be considered a more geometrically adaptive model that can reduce to the spherical case when the geometry of the problem allows this. An example of this property is given in Figure 1B comparing spherical and spheroidal harmonics on spherical and spheroidal geometry.

### 2.2 Fitting harmonic models to MEG data

The observed OPM data (***Y***) is an *n*_c_ (number of channels) by *n*_t_ (number of timepoints) matrix. It is composed of the sum of an assumed white noise component (***ε***) as well as interference (***E***) and lead fields (***L***) multiplied by some neural current (***J***), corrupted by multiplicative calibration and/or orientation errors (***C***) due to cross-talk (Nardelli et al., 2019), CAPE (Borna et al., 2022) and drifts (Iivanainen et al., 2019). That is, in the case of uncorrupted measurements, **C** would be the identity matrix.

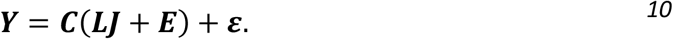

This data (***Y***) can be modelled as the linear combination (***β***) of harmonics (***H***) which can be spherical or spheroidal

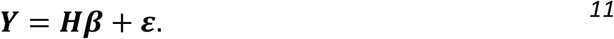

These harmonics (***H***) are partitioned column wise into harmonics that model both space inside the array (***H***_***in***_) and the space outside the array (***H***_***out***_)

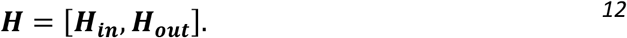

By inverting the harmonic model and multiplying this inverse (***H***^+^) by the data (***Y***)

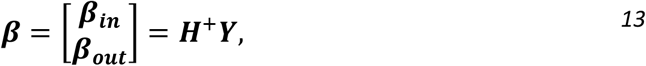

we then obtain the weights (***β***) which describe the relative contribution of the brain space (***β***_***in***_) and the interference space (***β***_***out***_). The data can then be replaced with a model of the data (***Ŷ*** _***in***_) which is constructed by multiplying the inner harmonics (***H***_***in***_) by the inner weights (***β***_***in***_)

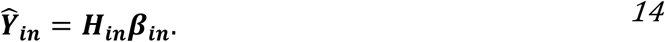

This model (***Ŷ*** _***in***_), when utilising spherical harmonics, is termed SSS in the literature (Taulu et al., 2005; Taulu & Kajola, 2005). The formulation presented above can be equivalently represented in a single matrix multiplication with an oblique projection matrix 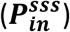 by utilising the fact that the parameter estimates (***β***_***in***_) of column wise partitioned matrices can be computed compactly (Baksalary & Baksalary, 2007)

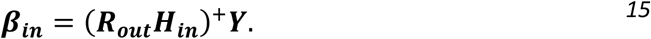

Where ***R***_***out***_ defines the projection orthogonal to the external space

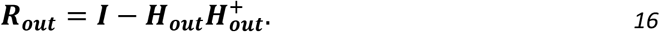

Now we can define the modelled data 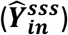 in a single oblique projection

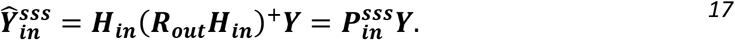

This projection is not guaranteed to be stable for all types of MEG systems (as it is an oblique projection) and can be very sensitive to white noise if the condition number (ratio of largest to smallest singular values) of the harmonic matrix (***H***) is high. Typically, in SSS this is addressed by optimising the harmonics in the basis set (Wang et al., 2023) or by optimising the system so that the resulting basis set will always be well conditioned (Zhdanov et al., 2023).

In contrast to these approaches, we propose the use of a projector 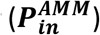 that expresses the neuronal data as the inner space signal components that are *orthogonal* to the interference space.

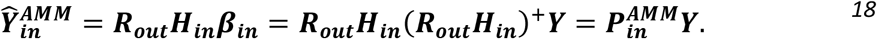

This is guaranteed to be a stable operation because the variance of data after an orthogonal projector is always less than the variance of the input data (see Appendix I). We should note that the AMM spatial projector does not explicitly fit a model of the exterior space as the inner space model is, by definition, orthogonal to exterior space. As such the degrees of freedom required to use AMM are solely determined by the order of the inner space. This means that AMM can be used with systems of lower channel count than SSS. Regardless of whether one uses the SSS style projector 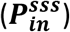 or the AMM style projector 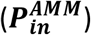 both are applied to the data (eq. 10) by a simple matrix multiplication

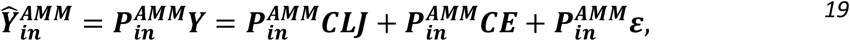

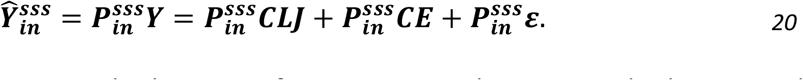

The motivation for summarizing this harmonic fitting into a single matrix multiplication is that now we can independently examine the effect of the type of projection on brain signal (***LJ***), white noise (***ε***) and interference (***E***), as the operation is a linear one.

#### Fitting temporal harmonic models to MEG data

Temporal subspace intersection has been proposed as an effective approach for mitigating interference due to calibration and truncation errors, which is not well captured by ***H***_***out***_ (Golub & Van Loan, 1996; Sekihara et al., 2016; Taulu & Hari, 2009; Taulu & Simola, 2006). This method relies on the definition of two additional spaces, an outer/interference space 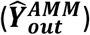 and an intermediate space 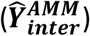. For AMM, the definitions of these spaces differ slightly to the SSS definition (Taulu & Simola, 2006) and are given below

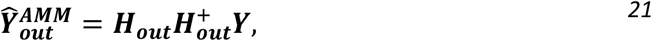

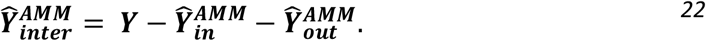

While the inner 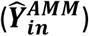 and outer 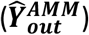 space contain the (spatial) brain signal and external interference respectively, the intermediate space 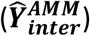 contains brain signals and interference signals that have a spatial order greater than that modelled by the inner or outer space. In other words, the intermediate space contains brain signals and interference that cannot be modelled with the inner and outer spaces. If, as a result of nonlinearity errors, there is residual interference in the inner space, there should exist a temporal correlation between the inner and *both* the outer and intermediate spaces. Furthermore, if one assumes that brain activity is not correlated over long time periods then this correlation will be purely due to interference.

In practice, this temporal correlation is calculated by finding the linear combination of channels in the inner and intermediate space (because it has more degrees of freedom than the outer space) that maximises the (canonical) correlation between these two spaces. If the correlation exceeds a certain limit (e.g 0.98), then the associated linear combination can be removed (under the assumption it reflects interference).

We provide the following heuristic (based on expected correlation of sinusoids in noise) for setting an upper bound on the correlation one can observe between inner and intermediate spaces. It incorporates previously empirically identified factors (Medvedovsky et al., 2009) such as the signal variance 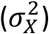, white noise variance 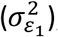, time window (*T*) and number of channels (*n*_c_). It also includes the expected fraction of neural signal in the inner (*α*) and intermediate spaces (*β*) and the maximum time window (*τ*) one would expect neural responses to be correlated over (e.g. 250 ms for an evoked response).

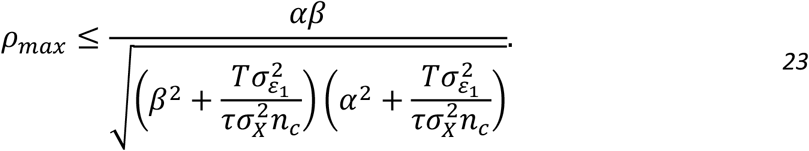

By setting a correlation limit at or above this threshold, one minimizes the risk of removing brain signal by chance. To give some insight into the intuition behind this heuristic, we provide an example in the supplementary material (Section S2) of using this bound to predict canonical correlations of sinusoids embedded in noise. In later sections we determine the scale factors (*α* and *β*) and use them to compare the bounds on worst case correlation between inner and intermediate spaces as a function of channel number and SNR. In practice, the above expression can be used to provide an adaptive threshold for a system with arbitrary numbers of channels and varying SNRs.

## 3. Methods

### 3.1 Lead field generation

OPMs were modelled as point magnetometers displaced 6.5 mm from the scalp. The source space was the MNI canonical cortical mesh (Mattout et al., 2007) available in SPM12 with 8196 vertices. The mean separation between vertices is approximately 5 mm. Lead fields were generated for each vertex of the source mesh. The orientation of each source was defined by the surface normal of the cortical mesh at that location. The forward model was the Nolte single shell model (Nolte, 2003). We investigate different sensor array types and layouts. Each array of sensors were placed on the scalp surface with inter-sensor separation ranging between 10 mm and 55 mm in steps of 5 mm. The sensor placement algorithm is described elsewhere (Tierney et al., 2020). For each level of sensor spacing we simulated single axis, dual axis and triaxial sensors, generating 3 lead field matrices per sensor spacing. Throughout the manuscript when we use the word “sensor” we are referring to a device that can potentially make multiple measurements in orthogonal directions. The word “channel” refers to 1 of these measurements (60 single-axis sensors have 60 channels; 60 dual axis sensors have 120 channels and 60 triaxial sensors have 180 channels). For dual axis sensing, the second axis was set orthogonal to the radial axis. To define this second axis we set the third component of the radial unit normal (say component r from vector [p,q,r]) to zero then swapped the remaining two components and negated the first (to give [-q,p,0]). This means the dot product of this vector with the original is zero (-pq+qp+0). The resulting vector was then normalised to have unit magnitude. For triaxial measurements, the third axis was defined by the cross product of the first two.

### 3.2 Effect of projector on neuronal space

In order to understand the order of spheroidal harmonic model required to sufficiently model the neural space, we multiplied both the SSS style projector 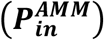 and the AMM style projector 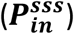 by the most densely sampled lead fields described in the previous section using both spherical and spheroidal harmonics. We then calculated the correlation (square root of variance explained) between the harmonic models and each and every lead field as a function of harmonic order (*l* = 1 to 11). We arbitrarily consider a minimum correlation of 0.95 across the whole brain as *‘sufficiently modelled*’. We repeated this projection and correlation for radial, dual-axis and triaxial sensing arrays. We set the order of the outer space to *l* = 2 throughout. Expected signal losses with higher or lower order external spaces is shown elsewhere (Tierney et al., 2022).

### 3.3 Effect of projector on interference space

It is well known that the ability of a multipole model to mitigate interface is strongly dependent on sensor linearity and the array design. This has been described elsewhere for SSS (Nurminen et al., 2008). As AMM is a fundamentally different projection (orthogonal vs oblique) the dependence on nonlinearity errors is not guaranteed to be the same. However, simulating realistic OPM nonlinearities is non-trivial as the calibration and orientation errors are a complex function of the magnitude of the interference and participant movement (Borna et al., 2022; Iivanainen et al., 2019). Rather than attempt to simulate realistic nonlinearities, we instead sought a nonlinearity topography that maximally disrupted our ability to reject interference. This may not be realistic but it provides a conservative lower bound on the effective shielding performance for the method.

To achieve this, we first simulated 500 interference topographies (composed of random combinations of homogenous field components and first order gradients). For each of these interference topographies we simulated 100 nonlinearity topographies that are also a random linear combination of homogenous field components and first order gradients. These nonlinearity topographies have a maximum magnitude varying between 0.5% and 10% of the interference topography magnitude. Thus, when the nonlinearity topography and the interference topography were multiplied (eqn. 10), a complex spatial dependency between the two was created. We then report the shielding factor from the resulting topography with the worst shielding factor. While this is not realistic, it represents a minimum expected performance that should generalise across environments and sensor designs. We repeated this process for radial, dual-axis and triaxial sensing arrays.

### 3.4 Effect of projector on white noise

The replacement of the OPM channel data by its harmonic components (either by AMM or SSS) will have an effect on the level of uncorrelated white noise over channels. This has recently been referred to as interpolation error or reconstruction error (Holmes et al., 2023; Zhdanov et al., 2023). As AMM is an orthogonal projection, we expect the white noise of the modelled data to always (regardless of system design) be lower than the original (or unprojected) data itself. In contrast, as SSS is an oblique projection, we expect the white noise to be a function of the system design and, in the cases of poor system design, greater than that of the original data. We applied both projectors (AMM and SSS) to Gaussian white noise vectors (0 mean, standard deviation of 1) with as many datapoints as we had channels for our simulated arrays (∼equally spaced on head surface with 55 to 10mm spacing in steps for 5mm). We calculated the white noise reduction factor as

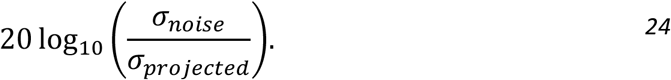

Where *σ*_*noise*_ represent the standard deviation of the white noise before projection and *σ*_*projected*_ represents the standard deviation of the noise after projection. We repeated this process for our radial, dual-axis and triaxial arrays.

### 3.5 Bounding correlation limits for temporal harmonic models

Having established the order of spheroidal harmonic (**section 3.1**) required to achieve at least a 0.95 correlation with the lead fields, we evaluated eqn. (23) as a function of channel number and SNR for a time window of 10s and an expected neuronal correlation time of 250ms. The intention here is not to be prescriptive in terms of correlation limits, but to demonstrate how one would theoretically select a correlation limit for a given SNR and array of a given channel count.

### 3.6 OP-MEG system description

To test this method empirically, a study was conducted at Young Epilepsy (Lingfeld, Surrey, UK), a charity for children and young adults with epilepsy. The shielded room, designed by Magnetic Shields Ltd (Staplehurst, UK), comprises two layers (outer and inner layers) of 1.5 mm thick MuMetal, and one layer (middle layer) of 4 mm-thick copper. An in depth description of the room, its active shielding and degaussing coils are described elsewhere (Holmes et al., 2022) and were utilised in the current study to minimise the background DC magnetic field. The OPM array used in this study was a 128-channel OPM array consisting of 64 QuSpin dual axis sensors provided by Cerca Magnetics Limited (Nottingham, UK).

### 3.7 Participant and paradigm

An 8-year-old, healthy male participated in this study. Informed consent was obtained and ethics were approved by the UCL research ethics committee. The participant passively viewed a reversing checkerboard (with check sizes subtending 5.4° degrees) projected on to the wall of the magnetically shielded room. The checkerboard reversed at a rate 4 reversals per second for a total of 200s resulting in 800 reversals. The reversals of the checkerboard were time locked to the OPM system by means of a photodiode.

### 3.8 Sensor level processing

Data was analysed using Statistical Parametric Mapping (SPM) (https://github.com/spm/spm) and the function for performing AMM (the combination of the spatial projector and the Canonical Correlation Analysis (CCA), section 2.3) is distributed within SPM (spm_opm_amm). Data were initially filtered between 1 and 70Hz using a 5^th^ order bi-directional Butterworth bandpass filter. Then both AMM and the SSS style projectors were applied to the data to create two separate datasets. The SSS projected data also had the temporal extension (TSSS) performed, as implemented in SPM. Both AMM and TSSS used a correlation limit of 0.98. The power spectral densities of all three datasets (filtered only, AMM and TSSS) were then calculated and shielding factors for AMM and TSSS were calculated relative to the filtered data. For the stimulus locked analysis, trials were subsequently epoched between 50ms pre-reversal and 1000ms post-reversal (to make 200 trials) and averaged across trials. T-statistics were also computed across trials to compare sensor SNR between filtered data, AMM projected data and the TSSS projected data.

### 3.9 Co-registration

An optical co-registration procedure was utilised to facilitate a source space analysis. An initial optical scan (https://www.einscan.com/handheld-3d-scanner/einscan-h/) was performed of the participant while they were not wearing the helmet. A second optical scan was performed while the participant was wearing the helmet. The template (canonical scalp) mesh available in SPM was then registered (using iterative closest point) to the individual scalp mesh (without helmet being worn). Subsequently these meshes were registered to the 3D optical scan mesh of the participant wearing the helmet using a 3 point fiducial match. Finally, these meshes were then registered (also using a 3 point fiducial match) to a 3D model of the OP-MEG helmet containing the sensor positions (provided by Cerca magnetics). At this point the template scalp and associated template brain are in the same space as the sensors and a source-level analysis is possible.

### 3.10 Source level processing

A standard minimum norm solution available in the Data Analysis in Source Space (DaiSS) toolbox (https://github.com/spm/spm/toolbox/DAiSS) was used to solve the inverse problem. For both TSSS and AMM, the lead fields were premultiplied by their respective, associated projectors. This lead-field correction is relatively uncommon for TSSS in cryogenic MEG settings. However, we will show later this is important for obtaining reasonable source localisation when applying TSSS to OPM data. The earliest detectable signal change post reversal (70ms) was localised using a t-statistic across trials. Lead field orientation was fixed in the direction of maximum SNR. The results are visualised on the MNI (Montreal Neurological Institute) template brain and thresholded at half maximum.

## 4. Results

### 4.1 Effect of projector on neuronal space

In Figure 2, the worst-case correlation (across all brain locations) between the harmonic model (spherical and spheroidal) and the lead fields is plotted as a function of harmonic order, using both the AMM and SSS style projector. We use the worst-case correlation as it provides us with the maximum number of channels that would be required to use this method. The correlation also indicates to what extent the model can reconstruct the data (with larger values being better). This is repeated for radial, dual axis and triaxial systems. We can see that for the SSS style projector (red lines), regardless of the number of channels per OPM in the system, the model converges towards a large correlation value (>0.95) as the harmonic order increases. It is interesting to note that the spheroidal model (squares) converges more quickly, meaning that fewer harmonics (and therefore channels) are required to model the data. Indeed, a correlation >0.95 is achieved for all systems using a harmonic order of 9. This means that when using spheroidal harmonics, *at most*, 99 channels (*l*^2^ + 2*l*) are required to fit such a model. Furthermore, we can see a key property of SSS here: SSS always preserves the brain signal components in a noiseless system.

**Figure 2.**
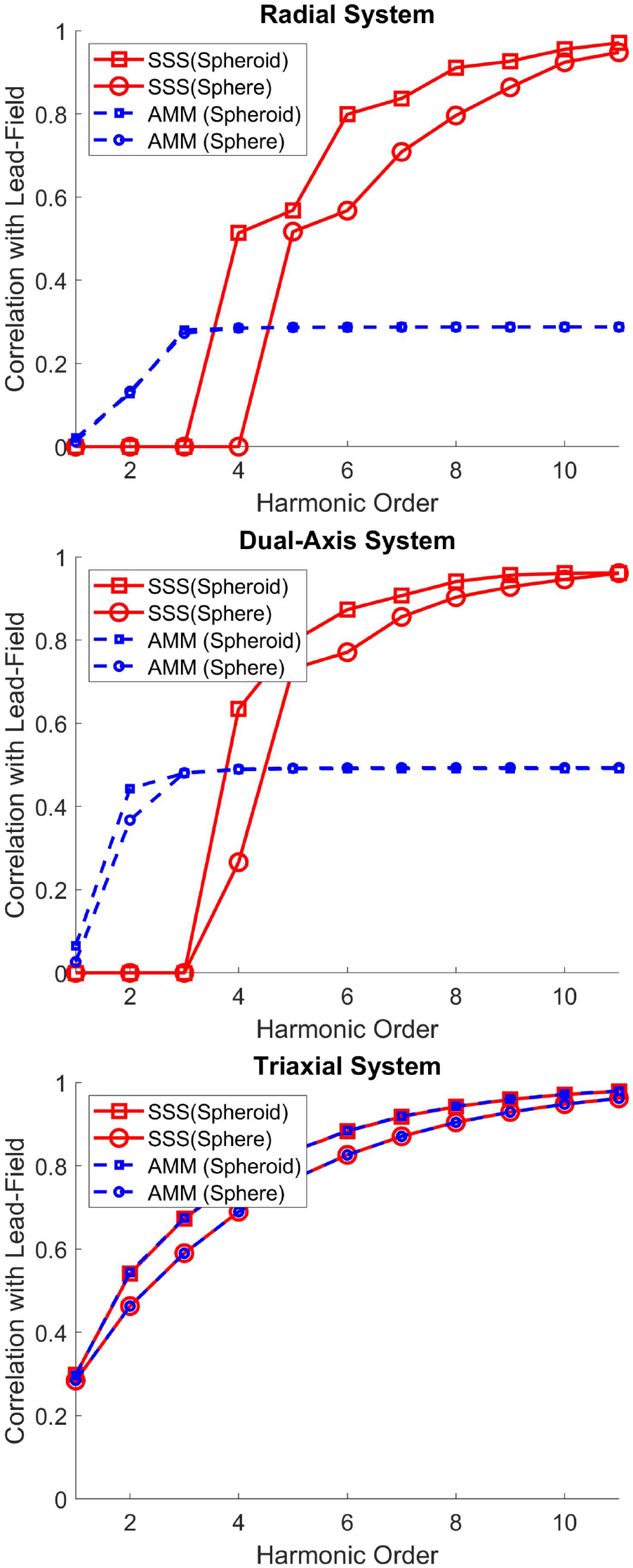
AMM and SSS projector effect on neuronal space. For all subplots, the lowest correlation across all brain vertices between the harmonic model and the corresponding lead field (Y axis) is plotted as a function of harmonic order (x-axis). The blue dotted lines depict the AMM style projector whereas the red solid lines depict the SSS style projector. The spherical and spheroidal harmonic models are depicted by circular and square symbols respectively. For all systems the SSS style projector reconstructs the brain signal with a worst-case correlation of 0.95 at order 9 for spheroidal harmonics order 11 for spherical harmonics. In contrast, the worst-case correlation for the AMM style projector is a function of system design and, at order 9, achieves a worst-case correlation of ∼0.333 for radial systems, 0.5 for dual axis systems and >0.95 for triaxial systems. While these correlations can be quite low (particularly for radial systems) it should be noted that by premultiplying the lead field by these projectors any distortion can be accounted for in subsequent modelling. The external order in all cases is 2.

In contrast, the AMM style projector risks removing brain signal components when utilising radial or dual axis systems. In the worst case, one could expect the correlation with the lead field to be reduced to a third for radial systems and to one half for dual axis systems. For triaxial systems both the AMM style projector and the SSS style projector produce near identical results (as interference and brain signal are approximately orthogonal) but the use of spheroidal harmonics means that lower order harmonics (and therefore fewer channels) can be used to model the neuronal space. We should note that while AMM can distort the topography of neural signals this distortion can always be built in to any forward model by premultiplying the lead fields by the same projector. This is common practice for methods such as signal space projection(Uusitalo & Ilmoniemi, 1997)

### 4.3 Effect of projector on interference

In Figure 3 the worst-case shielding factor is plotted for AMM and SSS as a function of the magnitude of spatially structured nonlinearity errors. Regardless of system, both methods share the same functional form for their dependency on nonlinearity errors (both are proportional to the reciprocal of the nonlinearity error). However, the AMM style projector has superior (and comparable) interference rejection for all systems and is more robust to the effect of spatially structured nonlinearity errors. For radial systems it is at least 3 times better, for dual axis systems it is at least twice as good, and for triaxial systems it is slightly better (10-15%). One can now clearly see that while using AMM can risk removing brain signal components (Figure 2) the trade-off is that it guarantees a minimum level of interference rejection across systems that is always better than or equal to SSS.

**Figure 3.**
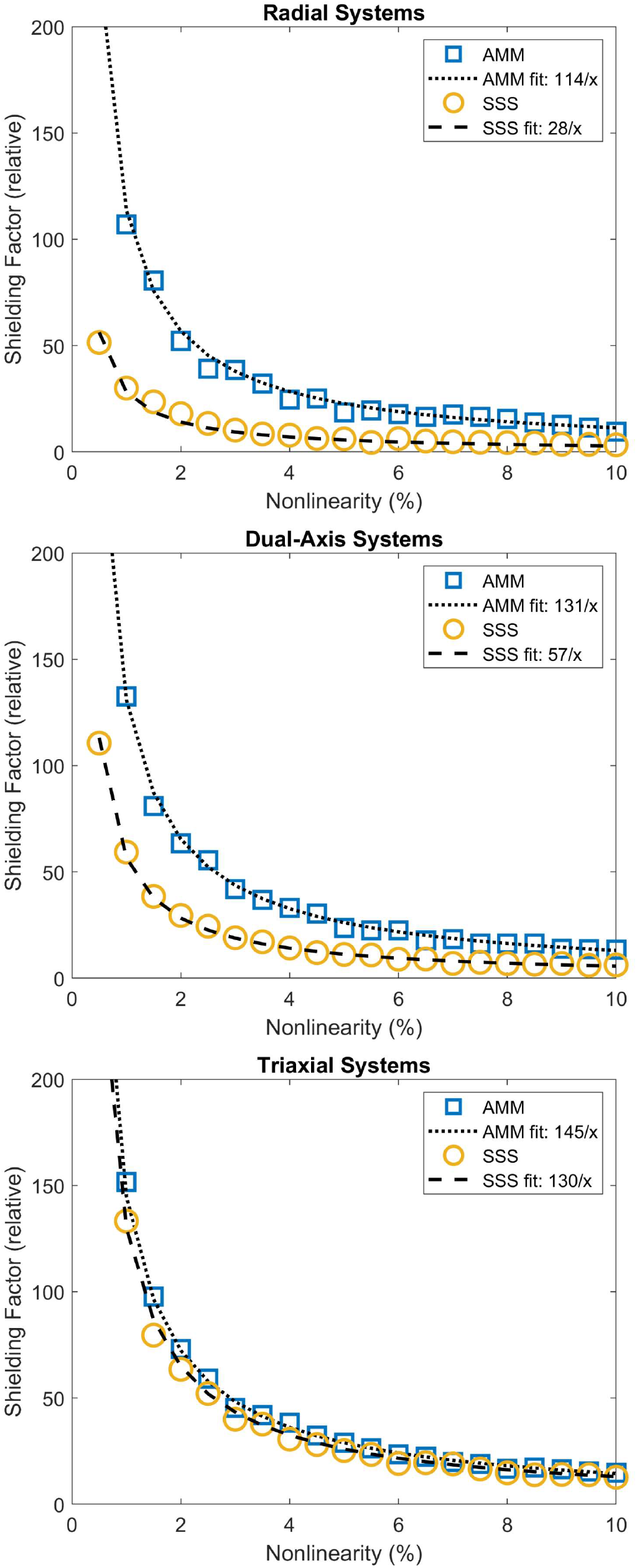
Worst-case interference mitigation in the presence of spatially structured nonlinearity errors. We simulated 500 interference topographies (composed of random combinations of homogenous field components and first order gradients). For each of these topographies we simulate 100 nonlinearity topographies that are also a random linear combination of homogenous field components and first order gradients. The nonlinearity topography and the interference topography are then multiplied, resulting in a complex spatial dependency between the two. We then report the lowest shielding factor (y-axis) obtained when applying the SSS style projector (yellow circles) and the AMM style projector (blue squares). We repeat this for each of the radial, dual-axis and triaxial systems as a function of the magnitude of the spatially structured nonlinearity errors (x-axis). We also report the line of best fit for this dependence (dotted lines for AMM and dashed lines for SSS).

### 4.3 Effect of projector on white noise

In Figure 4, we quantify the effects of the AMM and SSS style projectors on white noise. Regardless of the system, the AMM projector always reduces the white noise as the number of channels increases above the number of harmonics used to model the data. The relationship follows a simple square root law (e.g for twice as many channels as harmonics used in the model, the magnitude of the white noise magnitude is reduced by 3 dB, and for 4 times as many channels, 6dB reduction is achieved). This is because we are fitting more data to the same number of harmonic and orthogonal projections guarantee that the variance of the projected data must be less than the variance of the data itself.

**Figure 4.**
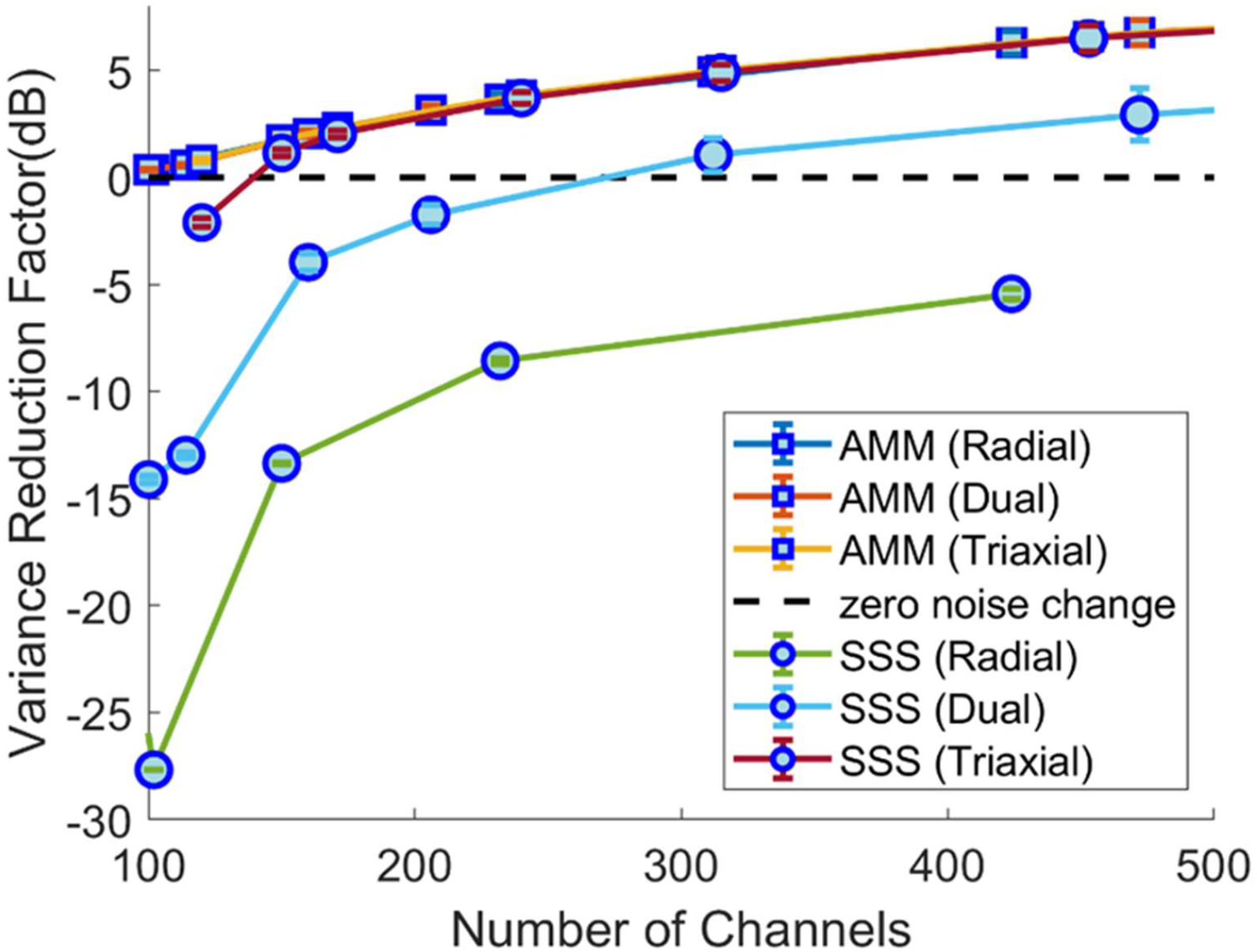
The effect of AMM and SSS projectors on white noise. The variance reduction factor in decibels (dB) (y-axis) is plotted as a funciton of channel count (x-axis) for both the the SSS style projectors (circles) and the AMM style projectors (squares). This is repeated for radial, dual and triaxial systems. The AMM projector always reduces the sensor white noise as it is an orthogonal projeciton (e.g reduced by 3dB at 200 channels and 6dB at 400 channels). However as the SSS projector is an oblique projection there is no guarantee on whether the variance will increase or decrease. Only in triaxial case does the SSS projector begin to reduce white noise effectively. This is because for triaxial systems the internal and external harmonics approach orthogonality and the oblique SSS projector approximates an orthogonal projection.

However, for the oblique SSS projector the same relationship is only observed for triaxial OPM systems with large channel count. This is because interference and brain signal become nearly orthogonal and thus the oblique projection of SSS approximates an orthogonal projection. The non-orthogonal nature of the SSS projection means that for both the dual axis and radial systems white noise will be increased for systems with channels counts less than 250 when using the SSS projector. For the on-scalp sensor arrays simulated here, there does not appear to be any reasonable channel count at which radial systems will not suffer from an increase in white noise.

### 4.4 Selecting correlation limits for temporal harmonic modelling

While the previous sections have focused on the spatial effects of AMM on OPM data, we now focus on how we can perform temporal harmonic modelling using CCA (similar to TSSS (Taulu & Simola, 2006) or Dual Signal Subspace Projection (DSSP) (Sekihara et al., 2016)) with OPM data of differing SNRs and channel counts. Figure 5 is an evaluation of eqn. 23 and is useful for setting a correlation limit (the highest temporal correlation that can exist between the neural and intermediate space by chance). Any canonical vectors with a correlation coefficient above this threshold can be removed from the data as they are unlikely to reflect brain signal. While channel count has a noticeable effect on this threshold, it is the SNR of the neural response that has the strongest effect on setting this limit. This is because even if a small percentage of a high SNR neural response leaks into the intermediate space it can have a high canonical correlation coefficient. By way of example, for single trial analysis, with SNR<10 and <200 channels, this figure suggests a correlation limit of 0.98 would be appropriate. For SNR<6 and <200 channels, a correlation limit of about 0.95 would be appropriate.

**Figure 5.**
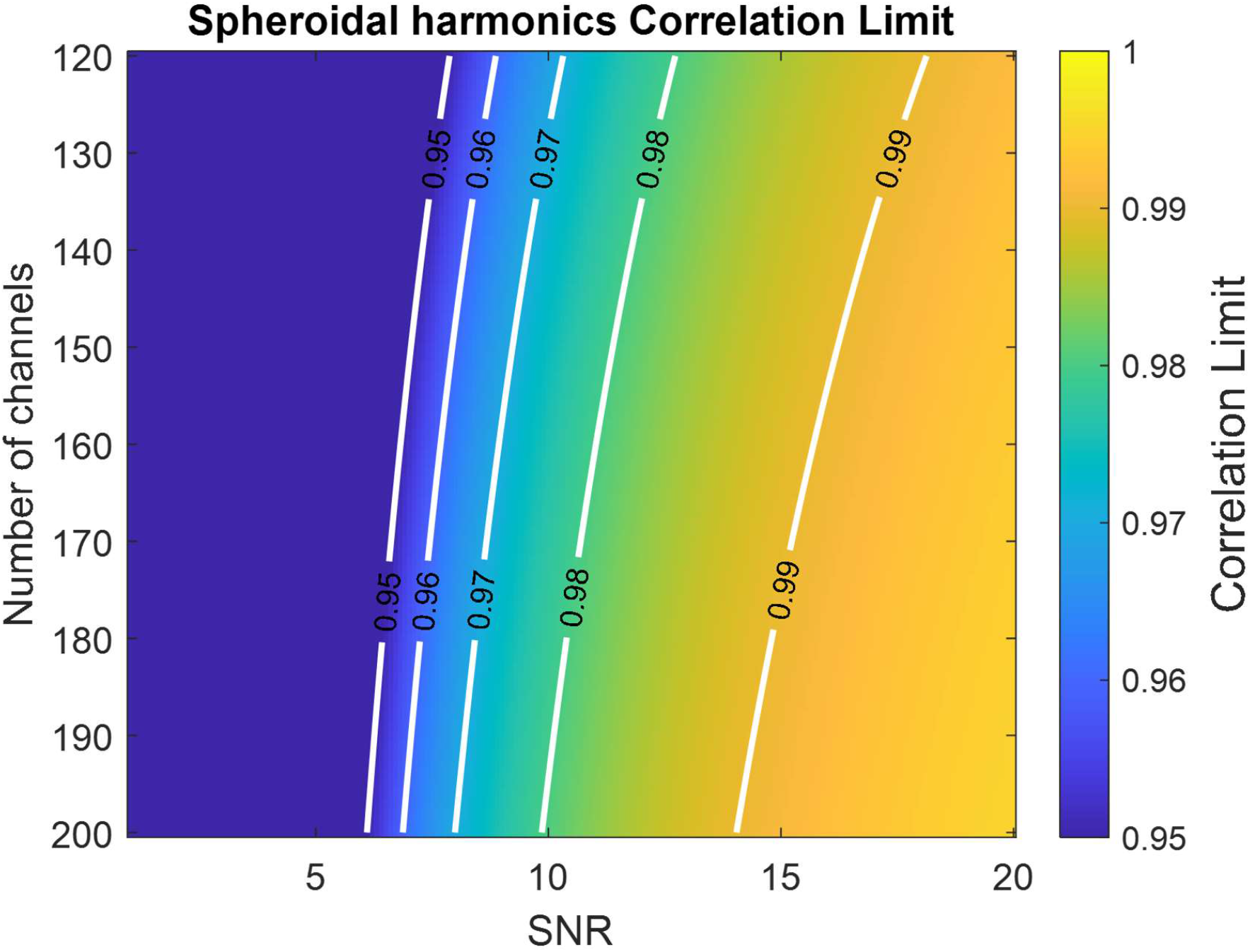
Setting corrleation limits for temporal harmonic analysis. The colour of the image reflects the maximum temporal canonical correlation coefficient that can exist between the neuronal and intermediate space. This limit is strongly influenced by SNR (x-axis) and channel count (y-axis). The order of the inner space is assumed to be order 9.

### 4.5 Power spectral densities

In Figure 6 we show the raw, filtered and processed (with AMM and TSSS) power spectral densities of the empirical, visual evoked potential data. We also show the shielding factors (relative to the filtered data). Both AMM and TSSS reduce the large low frequency interference and the 50Hz with AMM providing up to 6dB more attenuation (in this dataset). The most striking difference is that TSSS increases the broadband white noise substantially (in line with the simulation results). This effect is entirely driven by the oblique nature of the SSS spatial projection and how poorly conditioned the basis set spanning the neuronal and interference space is for this array. In contrast, for AMM there is minimal to no change in white noise across the spectrum. This issue can be mitigated for TSSS with array specific regularisation, but we found this caused a non-trivial temporal correlation between inner and intermediate spaces, thus complicating the temporal expansion of SSS, TSSS (see supplementary figure 2).

**Figure 6.**
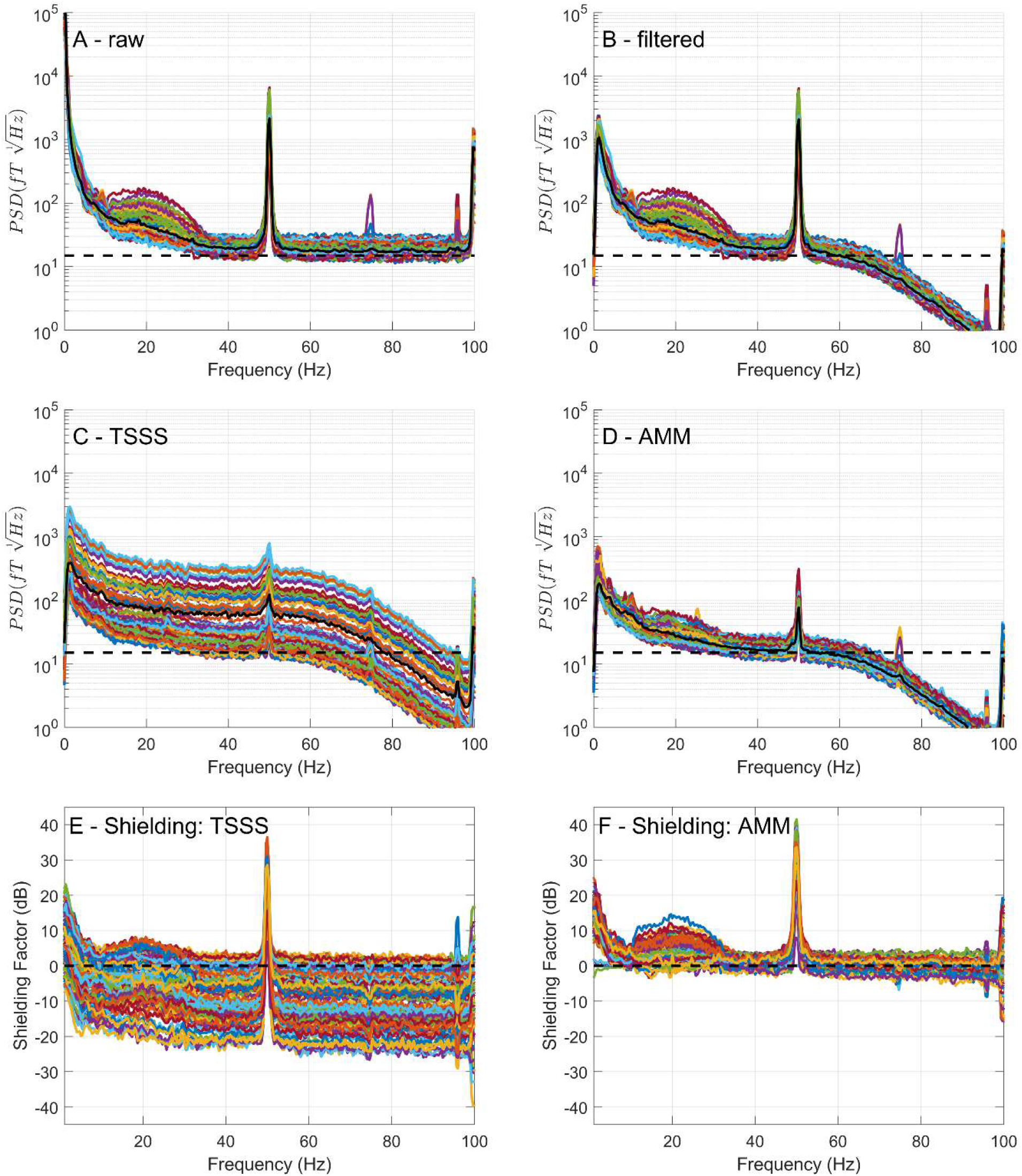
Power spectral densities. In the top four panels (A, B, C and D), the power sepctral densities (y-axis: rms) are shown for the raw, filtered, TSSS-processed data and AMM-processed data as a function of frequency. In the bottom two plots, the relative Power Spectrum Density (PSD)/software shielding factor (dB) is plotted as a function of frequency for AMM and TSSS relative to the filtered data. Both AMM and TSSS reduce the large low frequncy interference as well as the 50Hz, with AMM achieving up to 40dB of shielding. Where AMM and SSS markedly differ is the effect of the projectors on the broadband noise, which is increased in SSS. This is due to the poor conditioning of the basis set utilised and the oblique nature of the spatial projector (given the array used).

#### 4.5.2 Sensor Level results

While AMM has markedly reduced interference, without negatively impacting the white noise floor, it is possible that AMM can reduce the magnitude of the brain signal. As such, in Figure 7 we look at the sensor level results for the response to the checkerboard over a period of 1s. There are 4 checkerboard reversals per second and these 4 reversals are clearly visible in the AMM processed data (figure 7C). This response is visible in the raw data which has only been temporally filtered (figure 7A) but is corrupted by low frequency interference. In the TSSS processed data the instability (in the projection) obscures the response (figure 7B). In panels 7D-F the topography of the earliest component of the evoked response (70ms post reversal) across sensors is shown in terms of a t-statistic (or SNR). The spatial pattern is again similar for the temporally filtered (figure 7D) and AMM (figure 7F) processed data with the AMM data. The spatial pattern in the TSSS processed data is distorted to the extent a response is not clearly visible (we will show later that this distortion can be addressed in subsequent source modelling).

**Figure 7.**
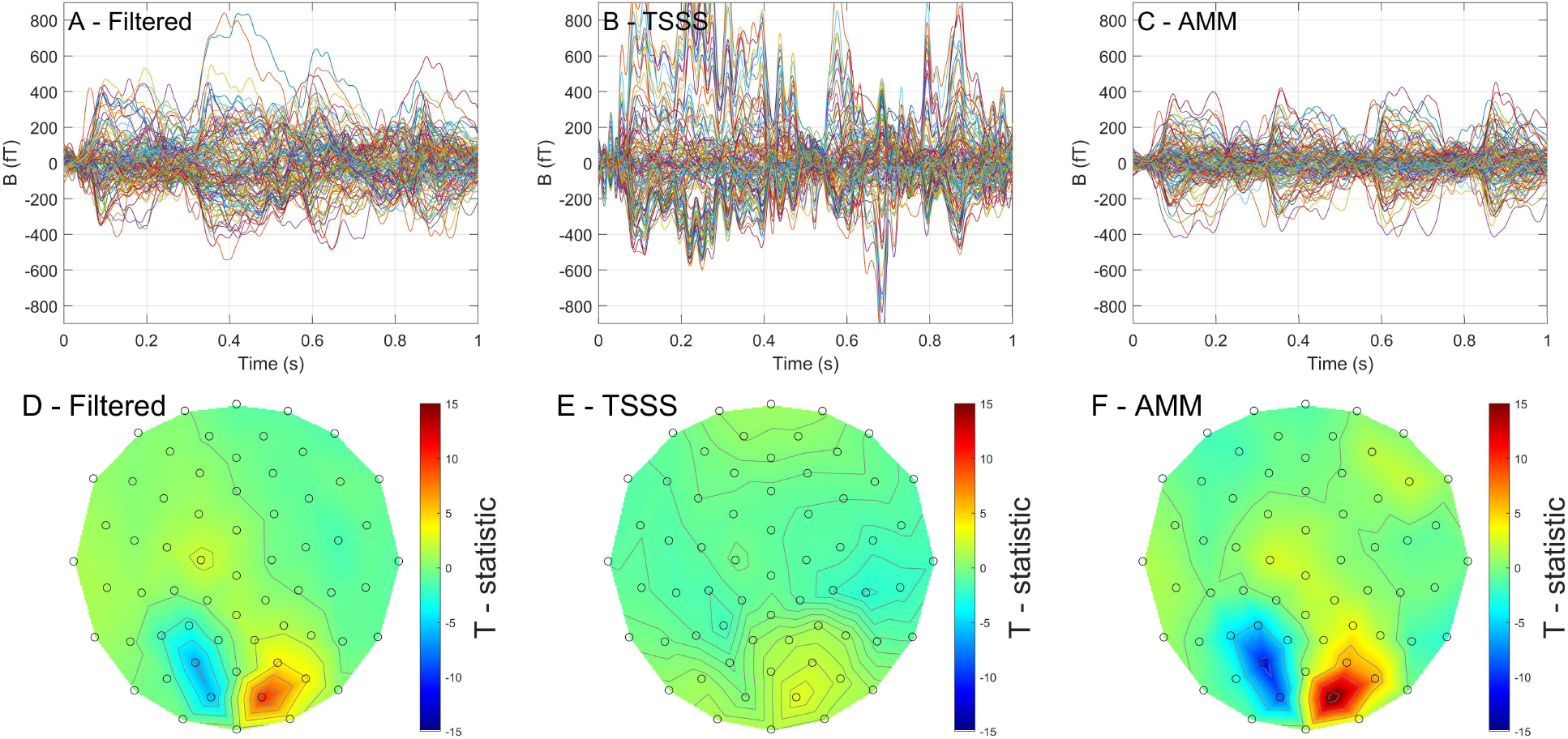
Sensor level visual evoked response. The top row shows the filtered data (A), TSSS processed data (B) and the AMM processed data (C) for all 128 channels. The data is epoched over a 1 second period (4 periods of checkerboard reversal, 200 averages). In the AMM processed data, the 4 responses are clearly visible but they are corrupted in the filtered data by low frequency interference. In contrast, for the TSSS processed data the poor conditioning of the basis set used results in a very unstable estimate of the evoked response (B). The bottom row shows the topography of the t-statistic (across trials) of the radial projection of the evoked response at 70ms. This effectivly quantifies SNR. The topography of the filtered (D) and AMM data (F) are quite coparable but the AMM processed data has better SNR (higher t-values). The topography of the TSSS (E) processed data is not quite interpretable due to the poor conditioning of the basis set.

#### 4.5.3 Source Level results

A common concern associated with pre-processing methods that reduce the spatial rank of the data is that they will have a negative impact on the source level analysis. Therefore, in Figure 8 we show a minimum norm inversion of the 70ms response (shown at the sensor level in the topographies in Figure 7). We find that all methods produce physiologically plausible maxima in the occipital lobe. However, both TSSS (2^nd^ row) and AMM (1^st^ row) have substantially higher SNR than the temporally filtered data (4^th^ row). It should be noted however that for TSSS to obtain this higher SNR the lead fields needed to be premultiplied by the associated projection matrix. If this is not done the SNR is lower (3^rd^ row) and is comparable to the SNR obtained from just filtering (4^th^ row).

**Figure 8.**
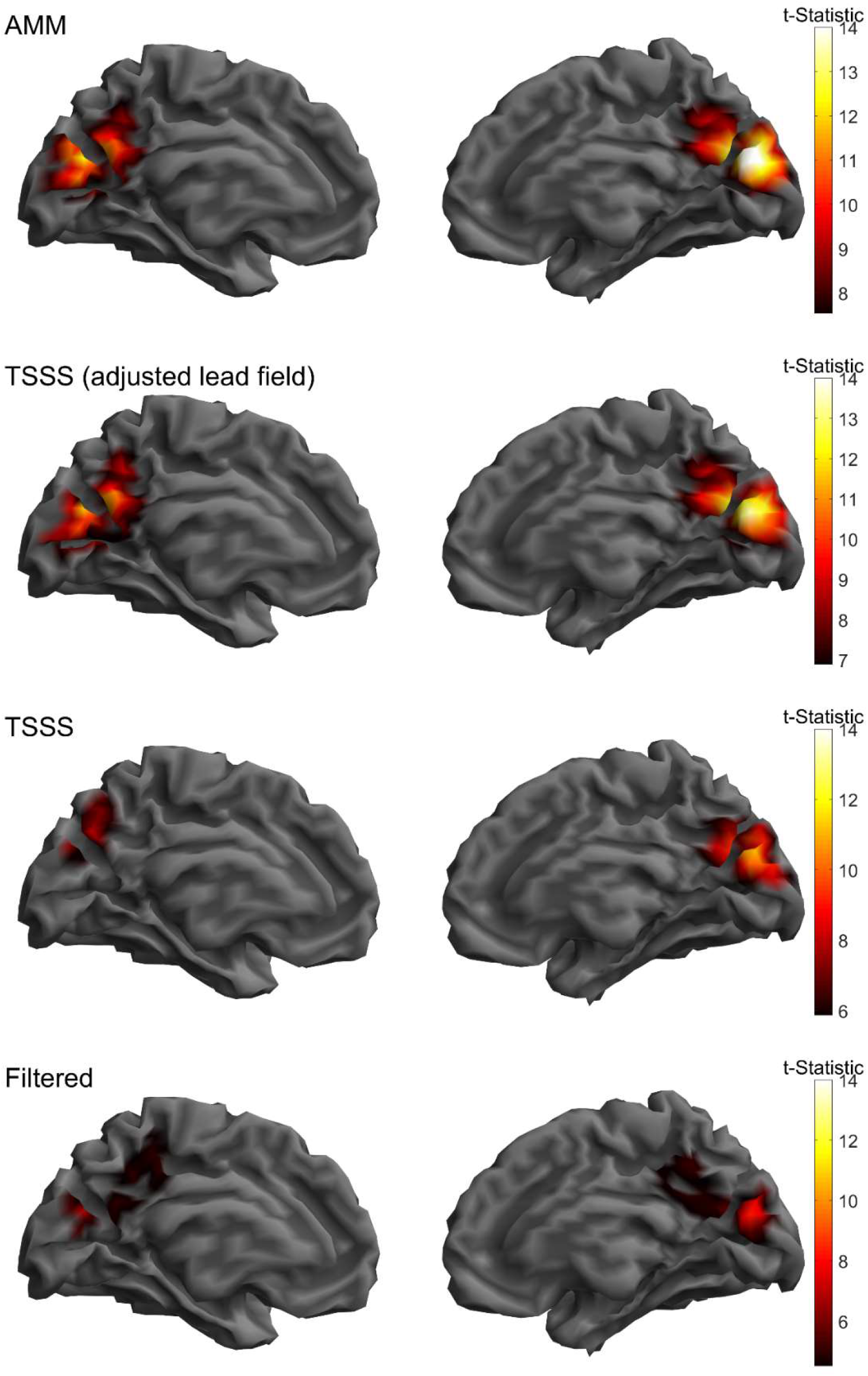
Minimum norm estimation of visual evoked repsonse. All rows plot the t-statisic (across trials) of the 70ms component of the evoked response on the template MNI brain mesh. All methods produce physiological maxima in the occipital lobe. However, AMM and TSSS produce better SNR than the purely tempoally filtered data. It should be noted that TSSS achieves highest SNR when the lead fields are premultipled (2^nd^ row) with the spatial projector (which is relatively uncommon for cryogenic MEG data). When the TSSS processed data does not have its lead fields premultipled by the projector (3rd row) the SNR is comparable to the filtered data (4^th^ row).

## Discussion

Here we have presented a method for modelling brain signal and interference, as observed by OPMs, as a linear combination of prolate spheroidal harmonics. These harmonics are fit to the data using a method that is stable for systems that are not optimally designed to separate brain signal from interference. We term this method AMM and verify its strengths and weaknesses in simulation. We also provide an empirical example of this method being used to model a noisy visual response to a flickering checkerboard at the sensor and source level. Finally, we have also contrasted the strengths and weaknesses of this method with the more commonly utilized multipole expansion, SSS.

In our previous work, we identified that using spherical harmonics to model brain signal, as observed by on-scalp magnetic sensors, was sub-optimal for brain regions near the front and back of the head (Tierney et al., 2022). This is because it is assumed when using spherical harmonics that all sensors exist outside a minimum volume reference sphere that encases the sources of interest. This condition can be easily violated in OPM data that is sampled on the scalp surface (Zhdanov et al., 2023). A similar problem is encountered in the field of gravitational modelling of asteroids. In such scenarios, it is common to model the gravitational potential using ellipsoidal or spheroidal harmonics (Fukushima, 2014; Hirt & Kuhn, 2017; Reimond & Baur, 2016; Winch, 1967). By applying these methods to simulated OPM data, we found that in the worst-case scenario the correlation between these harmonics and simulated brain signal never dropped below 0.95 across the whole brain when using order 9 harmonics (99 regressors). In contrast, to meet this same criterion using spherical harmonics required order 11 harmonics (143 regressors).

In addition to changing the basis set used for multipole modelling of OPM data, we also advocate to change how these harmonics are fit to the data. Typically, fitting these harmonics to data requires that the MEG system is designed so that the resulting basis sets have low condition number and therefore have a stable inverse. Due to the wide variety of OPM systems in development (Alem et al., 2023; Beato et al., 2018; Boto et al., 2022; Pratt et al., 2021) this is quite challenging. We find that poorly conditioned basis sets due to different system designs will result in poor interference rejection and increased white noise for the SSS style projector. However, when we use the AMM projector there is no increase in white noise and the shielding factors are greater than SSS even in the presence of spatially structured nonlinearity errors (regardless of system design).

In the worst-case scenario, AMM may reduce brain signal by up to two thirds for radial systems, one half for dual axis systems and negligibly for triaxial systems. This is in contrast to SSS which preserves the brain signal across all systems. These differences can be understood as the differences between orthogonal and oblique projections. The AMM only fits the portion of brain signal that is orthogonal to interference. It risks losing any brain signal that is spatially correlated with interference, but provides maximum interference rejection and stability with no arbitrary choice of regularization parameter. However, the oblique projections in SSS is fit to all brain signal, even if it is spatially correlated to interference. For systems that are designed sub-optimally, this results in increased uncertainty as to what is brain signal and what is interference resulting in decreased stability and poorer interference correction (in which case some kind of regularization choice is required).

In conjunction with the theoretical contributions and simulation results presented here, we also present an empirical example of using the AMM style projector and the SSS style projector to model OPM data obtained from a flickering checkerboard paradigm. We found, in line with the simulations, AMM to have higher maximum shielding factors and less sensitivity to white noise (Figure 6), leading to better SNR at the sensor level (Figure 7). Interestingly, we found, at the source level, TSSS and AMM produced comparable SNR if one pre-multiplies the lead fields by the SSS style projector (Figure 8). Effectively, if one performs the premultiplication of the lead fields by the projector, the forward model is aware of the oblique nature of the SSS projector and can account for it.

We should also caveat our source level results with a degree of uncertainty due to possible co-registration inaccuracies. We chose to use a generic helmet here, as opposed to the subject specific scanner casts we usually use (Boto et al., 2018; Meyer et al., 2017; Seymour et al., 2021; Tierney et al., 2018, 2018; Tierney et al., 2021). It is entirely possible that the helmet was moved (either via wobbling due to imperfect fit or by the participant themselves) from the position we obtained the 3D scan in. As such, the source level results should not be over-interpreted and more research regarding how harmonic models influence the breadth of source reconstruction methods is required.

We also stress that this work is not intended to be a direct comparison of any implementation of SSS (or TSSS, which were developed for use with cryogenic MEG systems) with AMM. Nor is the purpose of this work to describe how one may optimize SSS for a particular OPM system. That is now described elsewhere (Holmes et al., 2023; Nurminen et al., 2023; Wang et al., 2023; Zhdanov et al., 2023). As such, we did not explore the many ways in which the SSS method could have been stabilized. For instance, we could have explored how to optimally regularize SSS for OPM systems. We chose not to as the source space model could be stably estimated with good SNR when the lead fields were premultiplied by the SSS projector (although we do provide an example of the impacts of regularization on the sensor level PSD in supplementary Figure S2). We also could have excluded harmonics until the condition number reached an acceptable threshold (as can done in the SPM implementation for cryogenic MEG systems). We could have also selected the harmonics to be modelled using simulations that maximized SNR (Wang et al., 2023) or constructed the SSS design matrix in an iterative manner (Holmes et al., 2023). However, these approaches would need to be adapted for each and every system and our focus here is to provide a method that has more deterministic performance across a wide variety of OPM systems, with minimal custom modification.

Bearing these caveats in mind, we have developed a geometrically and statistically adaptive method of multipole modelling. We show in simulation and on real data that this model provides a compact representation of neuronal signal and interference, as observed by on-scalp MEG sensors. Shielding factors up to 40dB were observed in real data, even in the presence of spatially structured nonlinearity errors. The method’s inherent stability across systems, as it is an orthogonal projection, makes it an appealing preprocessing tool for the wide variety of OPM systems that are now becoming more readily available to the neuroimaging community.

## Appendix

### I: Stability of orthogonal projections

The squared norm of data (*Y*) premultiplied with a projector (*P*) is simply the inner product of the projected data with itself (*PY*)

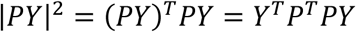

The key property of orthogonal projections that oblique projections do not have is that *P*^*T*^*P* = *P*. Therefore

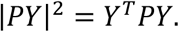

From the Cauchy schwarz inequality we know that the inner product of any two vectors is less than the product of their norms. As a result

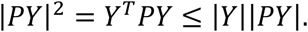

Cancelling the projected data norm (|*PY*|) from both sides we find that the norm (or variance) of the projected data (|*PY*|) is always less than that of the unprojected data (|*Y*|).

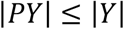

If we view algorithmic stability as the resulting changes in the output due to small changes In the input we can see that orthogonal projectors guarantee a stable output as the variance of the output will always be less than that of the input. This is not guaranteed for oblique projectors.

## Acknowledgements

SM was funded from an EPSRC-HIP award (EP/V047264/1). TT is funded by an ERUK and Young Epilepsy Fellowship (FY2101). The Wellcome Centre for Human Neuroimaging is supported by core funding from the Wellcome Trust (203147/Z/16/Z).

## Notes

### Competing Interest Statement

The authors have declared no competing interest.

