## Supplementary Material for "Adaptive multipole models of OPM data"

**S1: predicting canonical correlations**

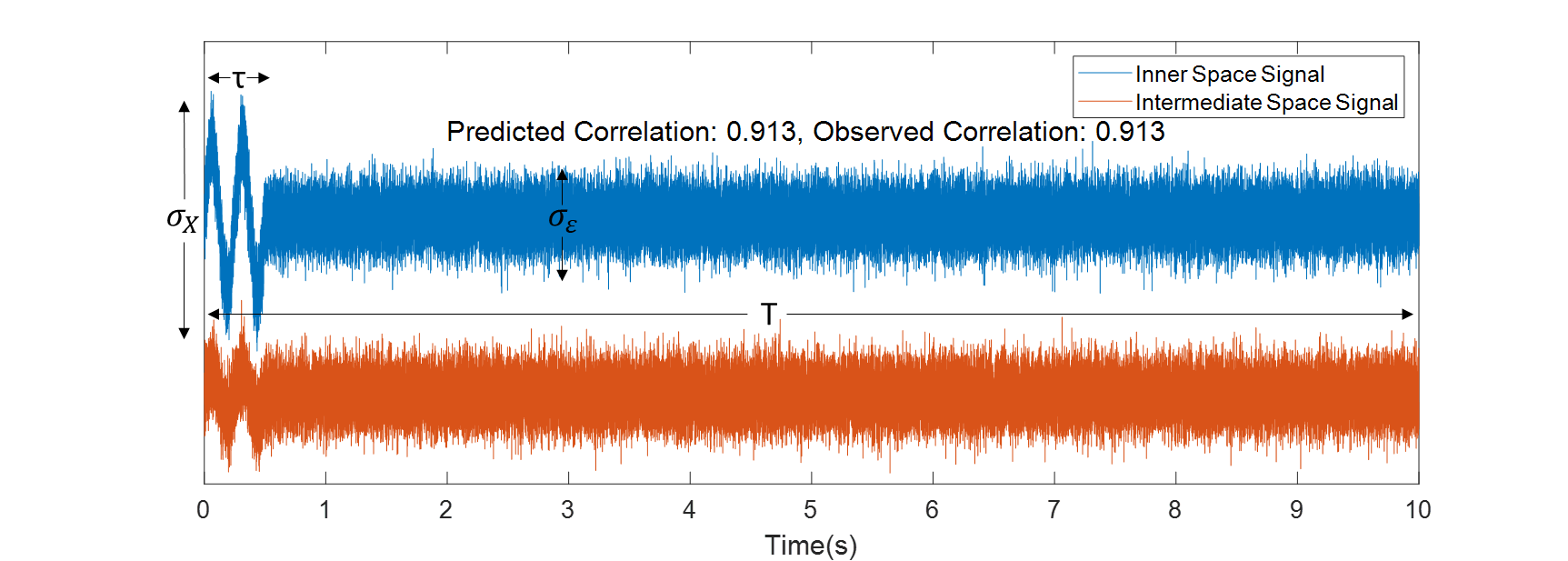

Figure S1. Predicting canonical correlation coefficients of sinusoids embedded in noise. Here we show a sinusoid, that is generated in the inner space (blue line) but due to truncation and calibration errors leaks into the intermediate space (red line). It has a particular standard deviation ($\sigma_{X}$) over a given time period ($\tau$). This signal is then embedded in white noise ($\sigma_{\varepsilon}$) over a longer time period ($T$). By making the unrealistic assumption that the magnitude of signal is the same across all channels (rather than only being present on a few) we can predict the maximal canonical correlation between the inner and intermediate space that can be generated by neural signals.

**S2: Regularising SSS**

In the main text we show that the parameters of the SSS model can be obtained by matrix inversion

$$\boldsymbol{\beta}=\left[ \begin{matrix} \boldsymbol{\beta}_{\boldsymbol{in}} \\ \boldsymbol{\beta}_{\boldsymbol{out}} \end{matrix} \right]=\boldsymbol{H}^{+}\boldsymbol{Y}.$$

However, this is not guaranteed to be a stable operation. We therefore explore the regularisation of this inversion ($\boldsymbol{H}_{\boldsymbol{reg}}^{\boldsymbol{+}}$) with the introduction of a parameter ($\lambda$)

$$\boldsymbol{H}_{\boldsymbol{reg}}^{\boldsymbol{+}}=\left( \boldsymbol{H}^{T}\boldsymbol{H}+\lambda\boldsymbol{I} \right)^{-1}\boldsymbol{H}^{T}$$

We vary lambda between 0 and 0.2 and find that with array specific regularisation the effects of SSS on broadband noise are diminished and approach that of AMM (Figure S2, left panel). However, the use of this regularisation parameter most likely results in real neural signal being assigned to the intermediate space (as it wasn’t classified as brain or interference). This caused exceptionally large temporal correlations to observed between the inner and intermediate spaces and when using a correlation limit of 0.98. As a result all signal was removed from the data using tSSS (Figure S2 B). It is clear that while array specific regularisation is possible for SSS using OPMs it creates a non-trivial coupling between the spatial projector and the temporal subspace intersection (tSSS). As AMM does not require any regularisation and our purpose here is not to optimise SSS or tSSS for OPM arrays we explore this issue no further.

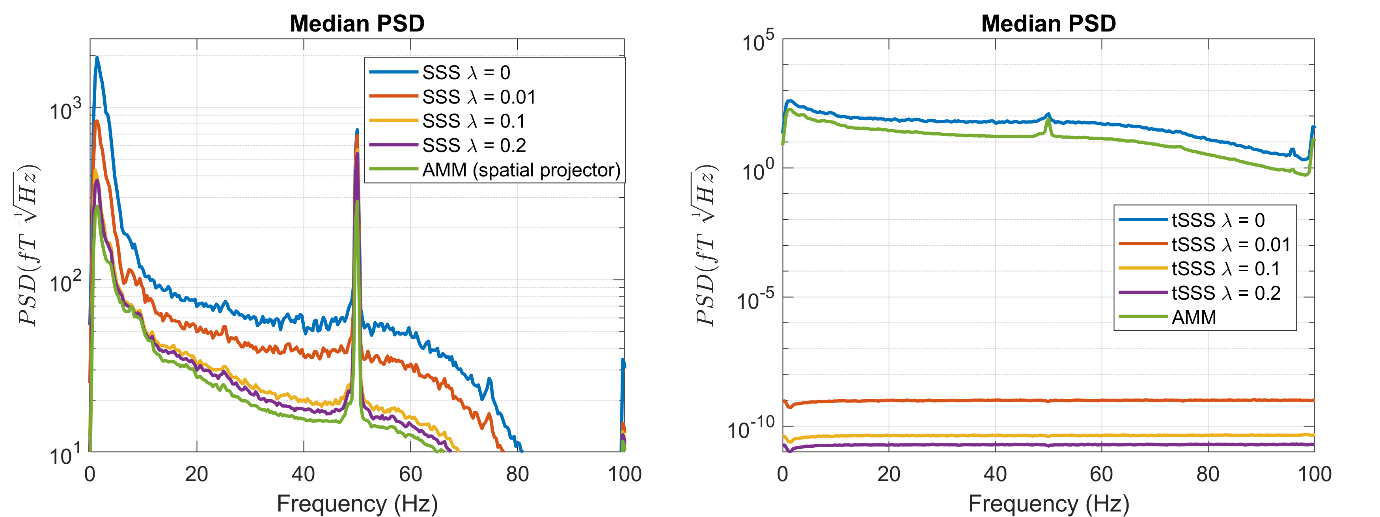

Figure S2. The effect of regularisation on SSS and tSSS. In the left panel we show how increasing the regularisation causes the SSS spatial projector to approach the performance of the AMM spatial projector. However, this parameter needs to be tuned each time an array is changed, whereas AMM requires no regularisation. Furthermore, in the right-hand panel, we show that using our default correlation limit (0.98) causes all signal to be removed when using tSSS that has had a regularised spatial projector applied. This is most likely because real brain signal has been assigned to the intermediate space, artificially inflating canonical correlations. As such, we have observed a non-trivial interaction between the regularisation parameter and the correlation limit that further complicates how one decides to regularise tSSS when applied to OPM data.
